# Cardiorespiratory fitness and cardiometabolic health are associated with distinct cognitive domains in cognitively-healthy older adults

**DOI:** 10.1101/2025.04.06.646611

**Authors:** Patrycja Kałamała, Nicholas Ware, Monica Fabiani, Patricia Michie, Montana Hunter, Alexandra Wade, Felicity Simpson, Maddison L. Mellow, Kathy Low, Hannah A.D. Keage, Gabriele Gratton, Ashleigh E. Smith, Frini Karayanidis

## Abstract

**Background:** Aging is associated with progressive cognitive decline, as well as increased prevalence of cardiometabolic risk factors and reduced cardiorespiratory fitness. In fact, reduced cardiometabolic health and cardiorespiratory fitness are both associated with a decline in cognitive functioning. This study examines the common and distinct contributions of these cardiovascular health factors on cognitive variability across different domains in cognitively healthy older adults.

**Methods and Results:** We apply structural equation modelling (SEM) to model cross-sectional relationships between cardiometabolic health, cardiorespiratory fitness and performance across multiple cognitive domains in an age-restricted sample of healthy older adults from the ACTIVate Study (n=345; 60-70 yrs). Participants completed a series of cognitive and clinical assessments (including brachial blood pressure, heart rate, blood-based metabolic markers). We designed a cognitive model (Model 1) with four latent factors that are differentially impacted by aging (Processing Speed, Executive Function, Verbal Memory and Crystallized Ability) and used it to test effects on cognition of two theory-driven dimensions of cardiovascular health: Cardiorespiratory Fitness (Model 2) and Cardiometabolic Health (Model 3). Model 4 included both predictors and examined their joint and distinct effects on these cognitive domains. When controlling for their joint variance, Cardiometabolic Health and Cardiorespiratory Fitness showed evidence consistent with a double dissociation on cognitive domains. Specifically, cardiorespiratory fitness significantly predicted processing speed (*r*=0.28, *p*<0.05) and executive function (*r*=0.66, *p*<0.05), but not verbal memory and crystallized ability. In contrast, cardiometabolic health predicted crystallized ability (*r*=0.31, *p*<0.05) and verbal memory (*r*=0.28, *p*<0.05), but not executive function and processing speed.

**Conclusions:** This study shows the first evidence that cardiorespiratory fitness and cardiometabolic health are associated with distinct cognitive domains in a large cross-sectional, age-restricted and high functioning cohort. These findings emphasize the importance of healthy aging approaches that target both health literacy and lifestyle behaviors to promote functional capacity across the lifespan.

## 1. INTRODUCTION

Aging is associated with structural and functional changes in the brain as well as a decline in a range of cognitive abilities (Caserta et al., 2009; Harada et al., 2013). Given the brain’s high energy demands, it is not surprising that these brain and cognitive changes are often associated with changes in cardiovascular health that become more prevalent amongst older adults. This paper examines the relationships between cardiovascular factors and cognitive health in community-dwelling older adults by modeling demographic, physiological, and cognitive variables from a large Australian older adult cohort (60-70 years).

Cardiovascular health is characterized by a range of physiological factors that can be used to estimate cardiometabolic health and cardiorespiratory fitness. Cardiometabolic health is informed by clinical markers, including conventional vascular risk factors (e.g., high blood pressure, abnormal lipid levels) as well as criteria relevant to metabolic syndrome (e.g., elevated waist circumference/BMI, high triglycerides; Cardiometabolic Risk Working Group, 2011). Cardiorespiratory fitness is a measure of functional capacity reflecting the integrated performance of the vascular, musculoskeletal, and respiratory systems (Kaminsky et al., 2019). It can be measured directly with cardiopulmonary exercise testing or estimated using validated predictors of VO₂max (Jurca et al., 2005). In this paper, we operationalize cardiometabolic health and cardiorespiratory fitness as latent constructs to represent these domains quantitatively.

Increasing age is associated with greater prevalence of cardiometabolic risk factors, such as hypertension, hypercholesterolemia, high blood glucose and obesity. These factors are associated with poorer cardiovascular health and associated diseases (Yaffe et al., 2020), as well as structural brain changes and greater risk of vascular dementia (De La Torre, 2012). Aging is also associated with a decline in cardiorespiratory fitness, which reflects the combined efficiency of the vascular, respiratory, and musculoskeletal systems (Kaminsky et al., 2019) and is linked to changes in brain structure and cognitive function (Bowie et al., 2021; Ross and Myers, 2023). Both cardiometabolic health and cardiorespiratory fitness are closely associated with efficiency of the cardiovascular system, tend to decline with increasing age and are associated with cognitive ability. However, their shared and distinct contributions to age-related cognitive decline are not well understood. With an increasingly aging population, it is essential to understand the distinct contributions of cardiometabolic health and cardiorespiratory fitness to cognitive functioning across different domains. This knowledge can inform the development of targeted, evidence-based interventions aimed at preserving cognitive function and enhancing quality of life in older adults.

### 1.1 Cardiometabolic Health & Aging

Cardiometabolic risk factors are highly prevalent even among otherwise healthy older adults, with 30% of older adults presenting with two or more such risk factors (Dove and Xu, 2023). In cross-sectional studies, the presence of cardiometabolic risk factors is associated with structural and functional brain changes, including greater cortical atrophy, white matter deterioration, and changes in cerebral blood flow regulation in response to neural activity (e.g., cerebrovascular reactivity) (Fuhrmann et al., 2019; Girouard and Iadecola, 2006; Kharabian Masouleh et al., 2018; Seshadri et al., 2004).

These cardiometabolic factors are also associated with variability in cognitive abilities, particularly in cognitive domains sensitive to aging, such as executive function, processing speed, and memory (Debette et al., 2011; Raz et al., 2008; Veldsman et al., 2020). Executive function^1^ regulates and coordinates cognitive resources to support goal-directed behavior and is highly sensitive to aging (Zanto and Gazzaley, 2017).

Processing speed, a key marker of psychomotor efficiency, also declines steadily with increasing age, contributing to broader cognitive slowing (Salthouse, 1996). Verbal memory (i.e., the ability to learn, retain and/or retrieve verbal information) declines to varying degrees depending on the specific task and context (Lamar et al., 2012; Olaya et al., 2017; Szoeke et al., 2016). In contrast, crystallized abilities (i.e., accumulated knowledge and vocabulary) remain relatively stable or even improve with increasing age (Lindenberger, 2001; Wang and Kaufman, 1993). This divergence between age-sensitive fluid abilities (e.g., executive function, processing speed, and verbal memory) and more stable crystallized abilities highlights the heterogeneous nature of cognitive aging and underscores the importance of investigating distinct cognitive abilities.

### 1.2 Cardiorespiratory Fitness & Aging

Cardiorespiratory fitness is a physiological measure that reflects an individual’s functional ability or their capacity to perform physical work. It relies on the ability of several physiologic systems to work synergistically to achieve and sustain physical activity (Kaminsky et al., 2019). Consequently, cardiorespiratory fitness is closely associated with measures of vascular, respiratory, and musculoskeletal function (Benck et al., 2017; Bhella et al., 2014; Earnest et al., 2013; Trappe et al., 2013). As these systems are involved in the delivery of oxygen (or the removal of metabolic byproducts) to the working organ, upstream deterioration (e.g., via aging or disease) in one or more of these systems reduces cardiorespiratory fitness (Kaminsky et al., 2019; Ross and Myers, 2023). In older adults, cardiorespiratory fitness is negatively associated with the prevalence and severity of cardiometabolic risk factors and the progression of cardiovascular disease, but positively with life expectancy (Mandsager et al., 2018; Snowden et al., 2013).

The rate at which cardiorespiratory fitness declines with increasing age is influenced by genetic predisposition (e.g., heritability estimates of ∼50% in sedentary individuals, (Bouchard et al., 1998)) and other determinants that affect respiratory, vascular, and musculoskeletal function (e.g., cardiometabolic risk factors, low physical activity, socio-economic status; Letnes et al., 2023; Loe et al., 2014; Zeiher et al., 2019). On average, cardiorespiratory fitness declines by approximately 8-10% per decade in healthy adults. By age 80 years, cardiorespiratory fitness may be reduced by as much as 50% relative to mid-life (Fleg et al., 2005; Forman et al., 2017).

While maximal exercise testing (e.g., VO₂max/peak) remains the gold standard for CRF assessment, non-exercise estimation models demonstrate high predictive validity (R² = 0.60–0.85) using readily obtainable variables such as age, sex, anthropometrics (BMI, waist-hip ratio), resting heart rate, and self-reported physical activity (Myers & Ross, 2021; Ross et al., 2016). These models provide a practical alternative to lab-based or submaximal field tests, enabling scalable CRF estimation in epidemiological and non-clinical settings.

Cardiorespiratory fitness is also associated with brain health and cognition during aging. Higher cardiorespiratory fitness is associated with better cognitive performance cross-sectionally, and the preservation of cognitive abilities longitudinally (Barnes et al., 2003; Bowie et al., 2021). Cognitive domains that are particularly vulnerable to aging (e.g., executive function, working memory, and processing speed) show the strongest associations with cardiorespiratory fitness (Erickson et al., 2022). Importantly, improving cardiorespiratory fitness may reverse age effects on cognitive and brain function (Colcombe et al., 2006; Dougherty et al., 2021; Freudenberger et al., 2016; Hopkins et al., 2012; Tyndall et al., 2013; Voss, 2010)

### 1.3 The current study

While both cardiometabolic health and cardiorespiratory fitness are closely linked to cardiovascular health and associated with age-related brain and cognitive changes, their common and distinct associations with cognitive outcomes are not well understood. In data from the Barcelona Brain Health Initiative (BBHI; *n* = 501, 53.6 years, 40-65 years), greater cardiometabolic risk was correlated with poorer performance across multiple cognitive domains, whereas cardiorespiratory fitness was more specifically associated with visuospatial and problem-solving abilities only in the older age range (55-65 years; (España-Irla et al., 2021). Whole brain cortical thickness mediated the relationship between cardiometabolic health and cognition, but cortical thickness in frontal regions more specifically mediated the link between cardiorespiratory fitness and cognition in the older subgroup. These findings suggest that cardiorespiratory fitness and cardiometabolic health may impact both common and distinct cognitive domains in middle-aged adults. However, the study did not investigate potential interactions between these two factors across various cognitive domains, which could provide a more nuanced understanding of their impact on cognition.

In the present study, we investigate the common and distinct associations of cardiorespiratory fitness and cardiometabolic health to four key cognitive domains in a large cohort of community-dwelling adults. The cohort was selected to be aged 60-70 years to minimize age-related variability. The cognitive domains were selected to represent both abilities that tend to remain largely unaffected with increasing age (i.e., crystallized abilities; (Tucker-Drob et al., 2022)) as well as abilities that show different degrees of decline (i.e., verbal memory, executive function, processing speed; Harada et al., 2013; Salthouse, 2010). Cardiometabolic health was operationalized as a latent variable using biological measures (i.e., systolic and diastolic blood pressure, glucose, high-density lipoprotein, triglycerides) and anthropometric indicators (i.e., BMI), consistent with established clinical risk assessments (Bitton and Gaziano, 2010; Goff et al., 2014; King et al., 2023). Cardiorespiratory fitness was operationalized as a separate latent variable using measures derived from established fitness estimators (Jurca et al., 2005; Wang et al., 2019), which included accelerometry measures of physical activity, heart rate, and anthropometric indicators (i.e., BMI).

Using structural equation modeling (SEM), we first created a model of cognitive ability with four latent cognitive factors covering the above domains (Figure 1, Model 1). Models 2 and 3 assessed the unique associations between these cognitive domains and cardiorespiratory fitness and cardiometabolic health, respectively. Model 4 included both measures of cardiovascular health in order to examine whether the pattern of unique associations found in Models 2 and 3 remain when controlling for their shared variance.

**Figure 1:**
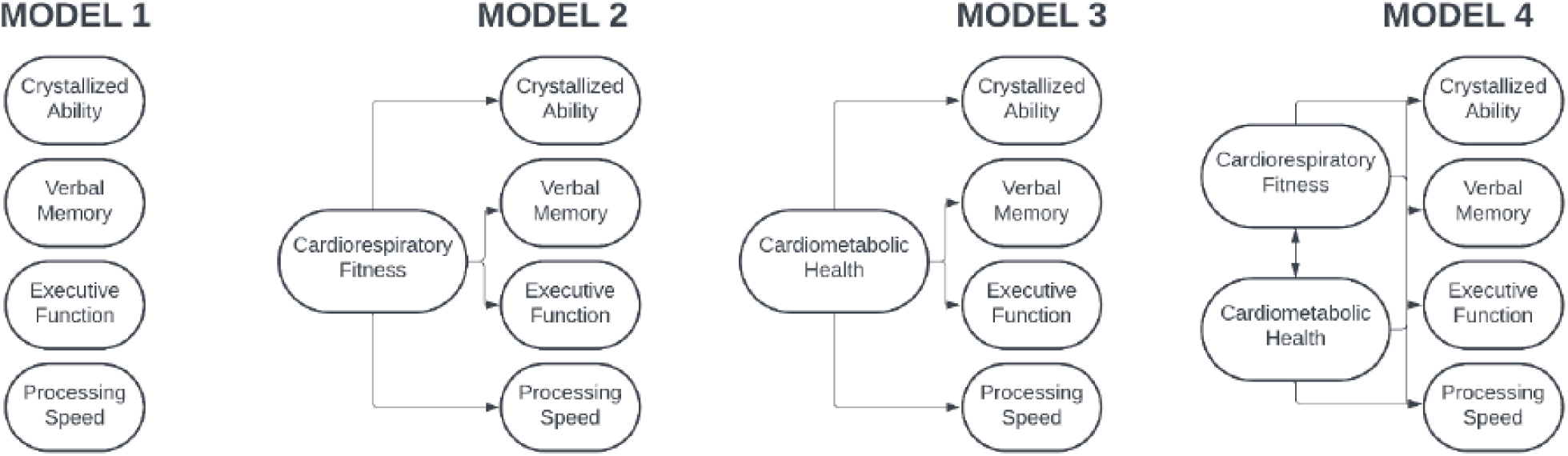
Schematic structure of conceptual models. Model 1: Four cognitive ability factors. Model 2: Direct associations between Cardiorespiratory Fitness factor and each cognitive ability factor. Model 3: Direct relationships between Cardiometabolic Health factor and cognitive ability factor. Model 4: Direct association between Cardiorespiratory Fitness and Cardiometabolic Health factors, and associations of each with cognitive ability factors, when controlling for common variance. Created in Lucid (lucid.co).

## 2. METHODS

### 2.1 Ethics

The current study includes Phase 1 data from the longitudinal ACTIVate Study (Smith et al., 2022) collected between August 2020 and February 2022 at the University of South Australia and University of Newcastle. ACTIVate was registered with the Australia New Zealand Clinical Trials Registry (ACTRN12619001659190) on November 27, 2019. Ethics approval was obtained from the University of South Australia and registered with the University of Newcastle Human Research Ethics Committee (H-2019-0421).

### 2.2 Participants

A total of 426 individuals met inclusion criteria for the ACTIVate Study (60-70 years, fluent in English). Participants were excluded if they had color-blindness, a clinical diagnosis of a major neurological or psychiatric disorder, intellectual disability or major physical disability, mild cognitive impairment or dementia (i.e., a score < 13/22 on the Montreal Cognitive Assessment – Blind/Telephone version; MoCA-B; Julayanont & Nasreddine, 2017), as assessed via telephone interview. Participants were primarily white Anglo-Saxon, with 79% born in Australia.

A total of 155 participants did not complete the task-switching paradigm and an additional 45 participants had missing data on physiological measures and/or other neuropsychological tasks, resulting in a complete data set of *n* = 226. Of these, three participants performed below chance level on the CANTAB Multitasking Task or task-switching paradigm and were excluded (*n* = 223). Of the 155 who did not complete task switching, 33 also had missing data on other cognitive and/or physiological measures. The remaining 122 were included in the modelling using FIML procedures to increase statistical power (see below).

Consequently, the analyses reported here are derived from a sample of 345 participants. To verify whether missing data affected the pattern of results, we also ran the analyses using the smaller sample with complete data (*n* = 223). The pattern of findings remained largely unchanged (see supplementary analyses: https://osf.io/pu8st/).

### 2.3 Demographic Measures and Cognitive Tasks

Demographic, past and current clinical history were collected during a brief interview. Socioeconomic status (SES) was quantified using postcode data to derive the Index of Relative Socio-economic Advantage and Disadvantage (IRSAD) from Australian Census data. IRSAD measures socio-economic advantage (e.g., high income, skilled occupations) and disadvantage (e.g., low income, unskilled occupations), with higher scores indicating areas with greater relative advantage (Australian Bureau of Statistics, 2021). Overall cognitive functioning against age norms was measured using Addenbrooke’s Cognitive Examination III, a brief, cognitive screening test that encompasses five cognitive domains (i.e., Attention, Memory, Fluency, Language, Visuospatial Ability; Mathuranath et al., 2000).

Cognitive measures included in the models were derived from the Multitasking Task, Reaction Time task and Verbal Recognition Memory task of the Cambridge Neuropsychological Test Automated Battery (CANTAB, Cambridge Cognition, 2023; Sahakian & Owen, 1992), the Oral Reading Recognition Task and Picture Vocabulary Task of the National Institute of Health Toolbox (NIH) Cognition Battery (NIHTB-Cog; Gershon et al., 2010; Hodes et al., 2013) and the cued-trials task-switching paradigm (Karayanidis and McKewen, 2021). The specific measures selected from each of the above tests are outlined in Table 1 (all tables are at end of manuscript).

**Table 1.** Overview of the cognitive variables.

| Measurement Tool | Variable Name | Operationalisation |
| --- | --- | --- |
| NIH Toolbox Cognitive Battery<sup>a</sup>:<br>**Oral Reading Recognition Test (ORRT)**<br>Expressive language | Oral Reading Recognition | The proportion of correctly pronounced letters and words. |
| NIH Toolbox Cognitive Battery<sup>a</sup>:<br>**Picture Vocabulary Test (PVT)**<br>Receptive language | Picture Vocabulary Recognition | The proportion of correctly selected pictures that best match the orally presented words. |
| CANTAB<sup>b</sup>:<br>**Reaction Time (RTI)**<br>Psychomotor speed | Simple RT | The sum of median reaction time (i.e., time from stimulus appearance to initiation of the response) and median movement time (i.e., time from response initiation to response completion). |
|  | Choice RT | The sum of median reaction time (i.e., time from stimulus appearance to initiation of the response) and median movement time (i.e., time from response initiation to response completion). |
| CANTAB<sup>b</sup>:<br>**Verbal Recognition Memory (VRM)**<br>Verbal episodic memory | Immediate Free Recall | The total number of correctly recalled target words during the immediate free recall phase. |
|  | Immediate Recognition | The total number of correctly recognised target words and correctly rejected distractor words in the immediate recognition phase. |
|  | Delayed Recognition | The total number of correctly recognised target words and correctly rejected distractor words in the delayed recognition phase. |
| CANTAB<sup>b</sup>:<br>**Multitasking Test (MTT)**<br>Task-set shifting and interference control | MTT Multitasking Cost (IES) | The difference in median reaction time between multitasking blocks (random sequence alternating between two rules: respond to the side on which the arrow appears vs. the direction the arrow points) and single-task blocks (respond to either side or direction across all trials). |
|  | MTT Congruency Cost (IES) | The difference in median reaction time between incongruent (e.g., arrow in left side but points right) and congruent (e.g., arrow in left side and points left) trials, averaged across single and multitasking blocks. |
| **Task-Switching Paradigm (TSWT)<sup>c</sup>**<br>Task-set shifting and interference control | TSWT Multitasking Cost (IES) | Also referred to as General Switch Cost: The difference in mean reaction time between mixed-task blocks (randomly alternating between letter and number classification tasks) and single-task blocks (repeating the same task across all trials), (averaged across incongruent and neutral trials). |
|  | TSWT Congruency Cost (IES) | The difference in mean reaction time between incongruent trials (i.e., target contains features from both cued and uncued tasks that are mapped to different responses) and neutral trials (i.e., irrelevant target feature is not mapped to any response) averaged across single-task and mixed-task blocks. |

Note. <sup>a</sup>NIH Toolbox® for Assessment of Neurological and Behavioral Function Administrator's Manual (2006-2020); <sup>b</sup> Cambridge Cognition Ltd. CANTABclipse Test Administration Guide (2012); <sup>c</sup> See Karayanidis and McKewen (2021)

The cued-trials task-switching paradigm (TSWT; Karayanidis & McKewen, 2021) was delivered using the parameters and data cleaning approach outlined in Whitson et al. (2014). Briefly, participants were trained on two simple classification tasks (i.e*., is number odd or even, is the letter a vowel or consonant*) to establish cue-target and target-response mappings. Each trial consisted of a cue (i.e., colored cross) that reliably indicated which task to perform on the subsequent target (C-T interval = 1000 msec). Targets consisted of two characters. One was from the cued task (e.g., a letter, if the letter task was cued). The other character was either a non-alphanumeric character (*neutral target, e.g., % or &* that was not mapped to any response) or an exemplar from the uncued task that was mapped to an incongruent response (*incongruent target,* e.g., if the letter task was cued, and the letter was mapped to a left-hand response, the number was mapped to a right-hand response). Participants completed the letter and number tasks either alone (single-task block) or in a random alternating sequence (mixed-task block). Median reaction time (RT) and average error rate were estimated separately for all trials in single-task blocks and in mixed-task blocks, and for all incongruent and all neutral trials (see Table 1). Trials associated with an incorrect response, or with RT that was too fast (i.e., < 200 msec) or too slow (i.e., RT >= mean RT(condition) + 3 x SD of RT(condition)) were excluded from RT estimates.

### 2.4 Physiological and Fitness Measures

Physiological measures (i.e., systolic and diastolic blood pressure, resting heart rate, and fasted blood-based metabolic markers) and anthropometric measures (i.e., body mass index calculated from height and weight) were obtained using standard procedures (see Smith et al., 2022 for detailed protocol).

Total time per day in moderate-to-vigorous physical activity (MVPA) was quantified from accelerometry data recorded over 7 consecutive days using a triaxial accelerometer (Axivity AX3) on the non-dominant wrist. Data were pre-processed using Open Movement GUI software (OmGUI) and analyzed with a custom MATLAB toolbox (COBRA; MATLAB R2018b; see Mellow et al., 2022 for analysis protocol).

Hypertension and hypercholesterolemia were defined using Australian guidelines (Australian Heart Foundation, 2023) and self-report. Hypertension was considered present if the participant had an average systolic blood pressure reading over >140 mmHg, reported having hypertension or using blood pressure medication. Hypercholesterolemia was considered present if they had a total cholesterol reading over 5.5 mmol/L (213 mg/dL), reported having or having received a diagnosis of high cholesterol by a health professional or reported using statin medication.

### 2.5 Data Preparation

The SEM analysis included socio-demographic variables (age, sex, years of completed education), hemodynamic variables (systolic and diastolic blood pressure in mmHg, resting heart rate as beats per minute), and blood metabolic variables (HDL, glucose, and triglycerides), as well as BMI as a proxy for body fat and MVPA as an indicator of physical activity level. It also included eleven cognitive variables listed in Table 1. Multitasking cost and congruency cost scores from both tasks were adjusted for accuracy (i.e., RT/*p*(correct)), yielding the inverse efficiency score (IES; Townsend & Ashby, 1978). IES captures both the speed and accuracy of responses and has a more normal distribution than raw RT-based measures. This is particularly important when testing older individuals, who often show a speed-accuracy trade-off (Salthouse, 2000).

All variables were adjusted, so that higher values for all variables indicate better condition/performance. To achieve this, values for blood pressure, heart rate, glucose, triglycerides, BMI, simple and choice RT, multitasking and congruency costs were reversed. All variables were screened for their distribution and subsequently scaled(i.e., standardized to have *M* = 0 and *SD* = 1; see below). .

### 2.6 Structural Equation Modeling (SEM)

We employed a SEM approach to first model the structure of cognitive domains and then added the two cardiovascular latent variables alone and together as predictors of these cognitive domains (Figure 1). Full information maximum likelihood was used to account for missing task-switching data for 122 participants in order to maximize sample size.

The variance of the latent variables was fixed to 1. Model fit was assessed using standard indices, including the root-mean-square error of approximation (RMSEA), standardised root-mean-square residual (SRMR), and comparative fit index (CFI). Conventionally, a good fit is indicated by RMSEA < 0.05, SRMR < 0.08 and CFI > 0.95 (Kline, 2016).

The data were analyzed and visualized in R - version 4.1.0 (R. Core Team, 2022), using the following packages: “tidyverse” (Wickham et al., 2019), “psych” (Revelle, 2007), “Hmisc” (Harrell, 2025), “lavaan” (Rosseel, 2012). The code is available in the project repository (https://osf.io/pu8st/). Requests for data access can be made through the ACTIVate study (Smith et al., 2022).

**Model 1:** We tested the feasibility of a **Cognitive Ability Model** with four cognitive factors that vary in the degree to which they are impacted by age (Harada et al., 2013). The *Crystallized Ability factor* loaded on measures associated verbal knowledge (Oral Reading Recognition and Picture Vocabulary Recognition from the NIH-Toolbox-Cb), and years of education which is often a proxy for cognitive reserve (Stern, 2002). The *Verbal Memory factor* loaded on Free Recall, Immediate and Delayed Recognition scores from the Verbal Recognition Memory test. The *Executive Function factor* loaded on Multitasking and Congruency Costs from both the Multitasking Task and the task-switching paradigm. The *Processing Speed factor* loaded on Simple and Choice RT).

All factors were allowed to correlate with each other. The residual errors of variables derived from the same tasks were correlated to account for shared task-specific variance (i.e., MTT Congruency Cost with MTT Multitasking Cost, TSWT Congruency Cost with TSWT Multitasking Cost). By allowing for correlated residuals, the model better reflects the underlying structure of the data and reduces the risk of misinterpreting task-specific or methodological influences as true relationships between variables. Since the Processing Speed factor consisted of only two variables, their loadings were constrained to be equal. This constraint ensures that both indicators equally represent the “Processing Speed” construct and addresses the potential issue of under-identification when a latent variable has few but highly correlated indicators (Crawford and Lamarre Jean, 2021).

**Model 2: Cardiorespiratory Fitness Model.** The *Cardiorespiratory Fitness latent variable* included resting heart rate, MVPA and BMI, consistent with variables used in non-exercise fitness estimation methods (e.g., Jurca score; Harber et al., 2024; Jurca et al., 2005). Model 2 examined whether the Cardiorespiratory Fitness factor can predict cognitive abilities in the Cognitive Ability Model.

**Model 3: Cardiometabolic Health Model.** Based on established algorithms for assessing general risk of cardiovascular diseases (e.g., Framingham Risk Score; D’Agostino et al., 2008), the *Cardiometabolic Health latent variable* was derived using key measures of cardiometabolic condition: systolic and diastolic blood pressure, glucose, HDL, triglycerides and BMI. Model 3 examines whether the Cardiometabolic Health factor can predict cognitive abilities in the Cognitive Ability Model. The residual errors of variables derived from the same measurements were correlated (i.e., Systolic Blood Pressure with Diastolic Blood Pressure and High-Density Lipoprotein Cholesterol with Triglycerides).

**Model 4:** The final **Cardiovascular Health Model** included **Cardiorespiratory Fitness** and **Cardiometabolic Health** as latent variables. These were modelled concurrently to (1) statistically adjust for their covariance and (2) isolate their independent associations with specific cognitive domains (e.g., episodic memory, processing speed).

Residual errors of variables sharing physiological measurement pathways were permitted to covary (e.g., systolic/diastolic blood pressure; HDL cholesterol/triglycerides) to account for shared biological variance beyond the latent factors.

Consistent with established cardiovascular health frameworks, BMI was retained as an indicator of *both* latent variables, reflecting its dual association with cardiorespiratory capacity (e.g., via aerobic efficiency) and cardiometabolic risk (e.g., via adiposity-related pathways). To evaluate whether this shared indicator introduced confounding, we conducted a sensitivity analysis (Model 4) excluding BMI from both latent constructs.^2^

#### Controlling for Age

As the cohort was selected to have a narrow age range (10 years), we did not anticipate significant age-related influences. To confirm this, we reran Models 1-3 with age allowed to correlate with the Cardiorespiratory Fitness and Cardio-Metabolic Health factors and predict cognitive factors. These models showed substantially poorer fit and, importantly, age did not correlate with the latent cardiovascular variables or account for the observed relationships (see: https://osf.io/pu8st/). Therefore, we report SEM analyses without age.

#### Controlling for Sex

While this paper is not focused on sex differences, we also examined whether the pattern of relationships between cardiovascular health patterns and cognitive outcomes varied across sex. Factor values for cardiovascular and cognitive variables were extracted from Model 4. First, independent t-tests were conducted to evaluate group differences in Cardiorespiratory Fitness and Cardiometabolic Health based on sex. Next, we used linear regression to investigate whether sex moderated the relationships between cardiovascular health dimensions and cognitive factors. Specifically, we fitted models in which the cardiovascular predictor (Cardiorespiratory Fitness or Cardiometabolic Health) was included alongside one of the aforementioned variables and their interaction term. These models were tested separately for each cognitive factor: Crystallized Ability, Verbal Memory, Executive Function, and Processing Speed.

## 3. RESULTS

### Cohort Characteristics

Table 2 presents summary statistics of socio-demographic, physiological, and cognitive measures for the entire sample of the cohort of 345 older adults and separated by sex^3^. The mean age was 65 yrs with a range of 60-70 years, by design. On average, the cohort was highly functioning (mean ACE-III = 95; Hsieh et al., 2013; Velayudhan et al., 2014), with females performing slightly better than men. No other sex differences were observed in basic demographics. On average, the cohort was highly educated, with most participants having completed post-secondary education. The mean IRSAD score of 1023 indicated a moderate socio-economic advantage, above the Australian mean of 1002.6 and the New South Wales mean of 1016.

**Table 2.** Mean and standard deviation for demographic, physiological (with clinical cut-off for concern), and cognitive measures.

| Variable | Full sample<br>*n*=345 | Females<br>*n* = 239 | Males<br>*n* = 106 | t-test<br>(2-sided) |
| --- | --- | --- | --- | --- |
| Age (yrs) | 65.46 (2.97) | 65.31 (2.84) | 65.80 (3.22) | $t(343) = 1.41, p = 0.16$ |
| Education (yrs) | 16.64 (3.24) | 16.48 (3.31) | 16.98 (3.06) | $t(343) = 1.32, p = 0.19$ |
| ACE-III (dementia considered < 82/100) | 95 (4) | 95 (3) | 94 (4) | **$t(343) = -2.20, p = 0.03$** |
| SES (New South Wales mean = 1016.0) | 1023.3 (71.05) | 1021.40 (70.80) | 1027.58 (71.78) | $t(320) = 0.72, p = 0.47$ |
| MVPA (recommended 30 mins/day) | 88.69 (46.85) | 82.34 (42.62) | 103.02 (52.68) | **$t(343) = 3.86, p < 0.001$** |
| BMI (obese >30 kg/m<sup>2</sup>) | 27.00 (5.20) | 26.84 (5.49) | 27.36 (4.50) | $t(343) = 0.86, p = 0.39$ |
| Glucose (normal < 150 mg/dL) | 92.07 (12.50) | 91.37 (12.91) | 93.65 (11.43) | $t(343) = 1.56, p = 0.12$ |
| HDL Cholesterol (normal > 60 mg/dL) | 68.43 (20.42) | 73.30 (19.98) | 57.46 (16.92) | **$t(343) = -7.11, p < 0.001$** |
| Triglycerides (normal <150 mg/dL) | 101.93 (49.48) | 100.29 (46.97) | 105.65 (54.80) | $t(343) = 0.93, p = 0.35$ |
| Resting Heart Rate (normal 60-100 bpm) | 66.49 (10.13) | 67.92 (9.50) | 63.27 (10.80) | **$t(343) = -4.02, p < 0.001$** |
| Brachial Systolic BP (normal <140 mmHg) | 134.36 (17.78) | 130.49 (16.96) | 143.09 (16.51) | **$t(343) = 6.42, p < 0.001$** |
| Brachial Diastolic BP (normal <80 mmHg) | 81.04 (10.14) | 80.23 (9.68) | 82.86 (10.95) | **$t(343) = 2.34, p = 0.03$** |
| NIH Toolbox Reading Recognition | 119.00 (13.43) | 118.87 (12.58) | 119.31 (15.24) | $t(343) = 0.28, p = 0.78$ |
| NIH Toolbox Picture Vocabulary | 113.13 (9.83) | 113.38 (9.95) | 112.57 (9.59) | $t(343) = -0.71, p = 0.48$ |
| VRM Immediate Recognition | 32.38 (2.53) | 32.36 (2.59) | 32.44 (2.39) | $t(343) = 0.30, p = 0.77$ |
| VRM Delayed Recognition | 31.51 (2.79) | 31.65 (2.71) | 31.19 (2.97) | $t(343) = -1.43, p = 0.16$ |
| VRM Free Recall | 5.88 (2.21) | 6.04 (2.23) | 5.52 (2.14) | $t(343) = -2.02, p = 0.05$ |
| MTT Multitasking Cost | 299.90 (184.56) | 313.85 (193.22) | 268.42 (159.79) | **$t(343) = -2.12, p = 0.04$** |
| MTT Congruency Cost | 137.76 (104.23) | 146.16 (110.94) | 118.82 (84.67) | **$t(343) = -2.26, p = 0.02$** |
| TSWT Multitasking Cost | 170.17 (151.28) | 170.95 (137.78) | 168.67 (175.44) | $t(221) = -0.11, p = 0.92$ |
| TSWT Congruency Cost | 154.77 (82.13) | 164.69 (81.54) | 135.85 (80.36) | **$t(221) = -2.54, p = 0.01$** |
| Simple RT (msec) | 571.81 (78.75) | 572.44 (77.43) | 570.40 (82.00) | $t(343) = -0.22, p = 0.83$ |
| Choice RT (msec) | 660.58 (78.78) | 661.45 (80.11) | 658.63 (76.03) | $t(343) = -0.31, p = 0.76$ |

*Note.* Addenbrooke's Cognitive Examination III: A measure of global cognitive function, scored out of 100; SES: Socio-economic status, derived from the Index of Relative Socio-economic Advantage and Disadvantage (ABS, 2021); MVPA: Moderate-to-Vigorous Physical Activity, measured in minutes per day. For detailed descriptions of CANTAB, NIH and task switching variables, refer to Table 1. Significant differences ( $p < 0.05$ ) between females and males are bolded. Unless otherwise noted, analyses were conducted with $N = 345$ . Due to missing data, sample sizes for specific comparisons vary: $n = 322$ for SES and $n = 223$ for TSWT.

Participants were highly active (average of 1.5hrs/day MVPA), with higher activity in men (Table 2). Yet, BMI was in the middle of the overweight range for both sexes. On average, measures of cardiovascular health were within the healthy range (Nelson et al., 2024). Men had significantly higher systolic and diastolic BP, but lower resting heart rate and lower HDL than females (Table 2). As shown in Table 3, 53.3% of individuals did not meet the criteria for a hypertension diagnosis. Of the remaining 46.7% with hypertension, 24.1% were on one or more antihypertensive medication. Overall, 67.5% of individuals had hypercholesteremia and, of these, 16.2% were on medication to manage their cholesterol.

**Table 3.**
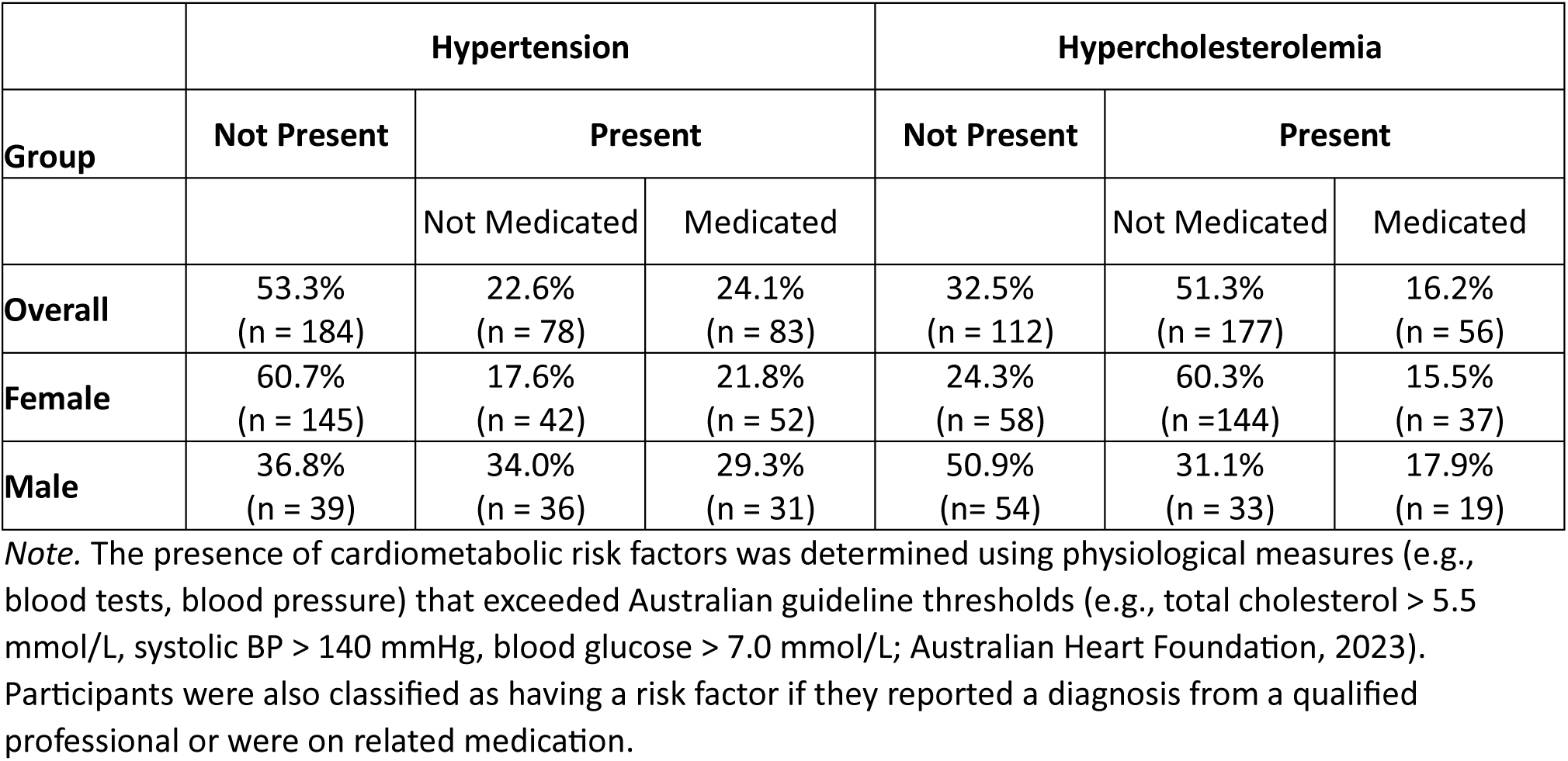
Presence of hypertension, hypercholesterolemia and medication status by sex.

|  | Hypertension |  |  | Hypercholesterolemia |  |  |
| --- | --- | --- | --- | --- | --- | --- |
| Group | Not Present | Present |  | Not Present | Present |  |
|  |  | Not Medicated | Medicated |  | Not Medicated | Medicated |
| **Overall** | 53.3%<br>(n = 184) | 22.6%<br>(n = 78) | 24.1%<br>(n = 83) | 32.5%<br>(n = 112) | 51.3%<br>(n = 177) | 16.2%<br>(n = 56) |
| **Female** | 60.7%<br>(n = 145) | 17.6%<br>(n = 42) | 21.8%<br>(n = 52) | 24.3%<br>(n = 58) | 60.3%<br>(n = 144) | 15.5%<br>(n = 37) |
| **Male** | 36.8%<br>(n = 39) | 34.0%<br>(n = 36) | 29.3%<br>(n = 31) | 50.9%<br>(n = 54) | 31.1%<br>(n = 33) | 17.9%<br>(n = 19) |

*Note.* The presence of cardiometabolic risk factors was determined using physiological measures (e.g., blood tests, blood pressure) that exceeded Australian guideline thresholds (e.g., total cholesterol > 5.5 mmol/L, systolic BP > 140 mmHg, blood glucose > 7.0 mmol/L; Australian Heart Foundation, 2023). Participants were also classified as having a risk factor if they reported a diagnosis from a qualified professional or were on related medication.

### 3.1 SEM Results

All variables exhibited substantial variability, as indicated by their standard deviations. Most variables were normally distributed, except for three metabolic variables (HDL, Glucose, and Triglycerides), which required log10 transformation to achieve normality.

Reliability of blood pressure and heart rate (three measurement points), BMI (two measurement points) were evaluated using the intra-class correlation coefficient. We did not assess the reliability of MVPA measures taken over seven days, as habitual activity varies across days and between weekdays and weekends. The reliability of measures derived from the task-switching paradigm was determined using split-half correlations and adjusted with the Spearman-Brown prophecy formula. As shown in Table 4, all measures were highly reliable. Reliability for the metabolic variables (i.e., Glucose, HDL, triglycerides) could not be established as we only had a single measurement. For CANTAB and NIH measures, we had limited access to raw data required for independent reliability calculations. Previous studies have reported adequate reliability for the NIH Toolbox and CANTAB tasks (see Gershon et al., 2013 for NIH; Karlsen et al., 2022; Skirrow et al., 2022 for CANTAB).

**Table 4.** Reliability of measurement tools.

| Variable | Reliability (r) |
| --- | --- |
| Systolic BP | 0.95 |
| Diastolic BP | 0.94 |
| Resting HR | 0.97 |
| BMI | 1.00 |
| TSWT Multitasking | 0.99 |
| TSWT Congruency | 0.99 |

*Note.* Intra-class correlation for systolic blood pressure, diastolic blood pressure, heart rate and BMI; odd-even Pearson's r with Spearman-Brown adjustment for TSWT Multitasking Cost and TSWT Congruency Cost. *N* = 223 for the TSWT Multitasking Cost and TSWT Congruency Cost correlations; *N* = 345 for the remaining.

Pearson correlations between key demographic and cognitive variables are reported in Table 5. As expected, variables related to cognitive abilities showed moderate-to-strong intercorrelations. Despite a restricted age range (60-70 years), measures contributing to the Processing Speed and Executive Function factors showed weak but significant correlations with age.

**Table 5:** Correlations between age and cognitive variables, age and physiological variables, cognitive and physiological variables.

| *A. Cognitive Variables* | 2<br>Educ | 3<br>RRec | 4<br>PVoc | 5<br>ImmR | 6<br>DelR | 7<br>FreeR | 8<br>MTT-M | 9<br>MTT-C | 10<br>TSWT-M | 11<br>TSWT-C | 12<br>sRT | 13<br>cRT |
| --- | --- | --- | --- | --- | --- | --- | --- | --- | --- | --- | --- | --- |
| Age | -0.02 | -0.01 | -0.02 | -0.02 | -0.04 | **-0.12** | -0.08 | **-0.21** | **-0.19** | **-0.19** | **-0.12** | **-0.16** |
| Education | — | **0.27** | **0.24** | 0.04 | 0.09 | 0.09 | 0.03 | **0.16** | **0.13** | **0.03** | -0.02 | -0.03 |
| Read Recognition |  | — | **0.47** | 0.09 | **0.23** | **0.25** | **0.15** | **0.19** | **0.24** | **0.17** | -0.08 | -0.10 |
| Pict Vocabulary |  |  | — | 0.10 | **0.14** | **0.19** | **0.13** | **0.15** | **0.25** | **0.12** | -0.06 | -0.05 |
| Imm Recognition |  |  |  | — | **0.62** | **0.40** | **0.12** | 0.05 | 0.06 | 0.05 | 0.06 | 0.05 |
| Delay Recognition |  |  |  |  | — | **0.46** | **0.16** | 0.10 | 0.06 | 0.10 | 0.05 | 0.02 |
| Free Recall |  |  |  |  |  | — | 0.00 | 0.06 | 0.10 | **0.16** | 0.05 | 0.02 |
| MTT Multitasking |  |  |  |  |  |  | — | **0.25** | **0.22** | **0.22** | **-0.12** | **-0.13** |
| MTT Congruency |  |  |  |  |  |  |  | — | **0.18** | **0.28** | 0.02 | 0.02 |
| TSWT Multitasking |  |  |  |  |  |  |  |  | — | **0.64** | 0.04 | 0.05 |
| TSWT Congruency |  |  |  |  |  |  |  |  |  | — | 0.09 | **0.14** |
| Simple RT |  |  |  |  |  |  |  |  |  |  | — | **0.84** |
| Choice RT |  |  |  |  |  |  |  |  |  |  |  | — |

| *B. Physiological Variables* | 2<br>MVPA | 3<br>BMI | 4<br>Gluc | 5<br>HDL | 6<br>Trigl | 7<br>rHR | 8<br>sBP | 9<br>bBP |
| --- | --- | --- | --- | --- | --- | --- | --- | --- |
| Age | -0.04 | -0.02 | -0.05 | -0.01 | **0.04** | -0.03 | **-0.14** | **-0.01** |
| MVPA (mins/week) | — | **0.36** | 0.04 | **0.20** | **0.26** | **0.21** | 0.02 | 0.02 |
| BMI (kg/m<sup>2</sup>) |  | — | **0.32** | **0.42** | **0.38** | **0.18** | **0.18** | **0.21** |
| Glucose (mg/dl) |  |  | — | **0.16** | **0.23** | **0.12** | 0.02 | **0.21** |
| HDL Cholesterol (mg/dl) |  |  |  | — | **0.48** | 0.04 | **0.12** | 0.06 |
| Triglycerides (mg/dl) |  |  |  |  | — | **0.14** | 0.09 | **0.17** |
| Resting Heart Rate (bpm) |  |  |  |  |  | — | -0.09 | **0.18** |
| Brachial Systolic BP (mmHg) |  |  |  |  |  |  | — | **0.62** |
| Brachial Diastolic BP (mmHg) |  |  |  |  |  |  |  | — |

| *C. Cognitive and Physiological* | MVPA | BMI | Gluc | HDL | Trigl | rHR | sBP | dBP |
| --- | --- | --- | --- | --- | --- | --- | --- | --- |
| Read Recognition | -0.07 | 0.08 | **0.13** | 0.05 | 0.04 | 0.04 | 0.07 | **0.16** |
| Pict Vocabulary | -0.00 | 0.08 | -0.02 | 0.05 | 0.00 | -0.03 | **0.14** | 0.05 |
| Imm Recognition | 0.05 | 0.02 | 0.00 | -0.00 | 0.07 | -0.04 | -0.01 | **0.11** |
| Delay Recognition | 0.01 | 0.08 | **0.11** | 0.04 | 0.07 | -0.03 | -0.04 | 0.09 |
| Free Recall | -0.00 | 0.09 | 0.07 | 0.03 | 0.05 | -0.05 | 0.04 | 0.10 |
| MTT Multitasking | -0.04 | -0.05 | -0.03 | 0.05 | 0.00 | **-0.12** | 0.00 | -0.01 |
| MTT Congruency | -0.03 | -0.05 | -0.11 | 0.07 | -0.10 | **-0.12** | -0.02 | -0.04 |
| TSWT Multitasking | **-0.13** | -0.07 | -0.04 | -0.05 | -0.07 | **-0.13** | -0.03 | 0.03 |
| TSWT Congruency | **-0.24** | -0.13 | -0.07 | -0.07 | **-0.15** | **-0.27** | 0.03 | 0.05 |
| Simple RT | -0.12 | -0.08 | 0.04 | -0.01 | -0.04 | -0.08 | 0.00 | -0.04 |
| Choice RT | **-0.14** | **-0.13** | -0.00 | -0.07 | **-0.12** | **-0.12** | 0.02 | -0.01 |

Note: For detailed descriptions of CANTAB, NIH and task switching variables, refer to Table 1. *p* < 0.05 bolded

Fit statistics show that all models demonstrated good-to-excellent fit (Table 6) and are described further below. All variables and factor scores have been adapted so that higher values represent better condition/performance.

**Table 6.** Goodness of fit statistics for the main models.

| Model | CFI | RMSEA [90% CI] | SRMR |
| --- | --- | --- | --- |
| Cognitive Ability | 0.981 | 0.034 [0.010, 0.053] | 0.041 |
| Cardiorespiratory Fitness | 0.974 | 0.033 [0.014, 0.047] | 0.044 |
| Cardiometabolic Health | 0.966 | 0.034 [0.021, 0.046] | 0.043 |
| Cardiovascular Health | 0.959 | 0.035 [0.024, 0.048] | 0.045 |
| Cardiovascular Health (no BMI) | 0.963 | 0.034 [0.022, 0.045] | 0.045 |

Note: CFI, Comparative Fit Index; RMSEA, Root Mean Square Error of Approximation; SRMR, Standardized Root Mean Squared Residual.

It is important to recall that variables contributing to the models below are scored so that high score represents better performance or condition.

#### Model 1: Cognitive Ability (Figure 2)

All factors significantly loaded onto their respective variables with coefficients varying between 0.36 and 0.92. For the Crystallized Ability factor, the two semantic knowledge tasks had higher loadings than years of education. For the Verbal Memory factor, both recognition scores had higher loading than immediate recall. For the Executive Function factor, all four cost variables had roughly equal loadings.

**Figure 2.**
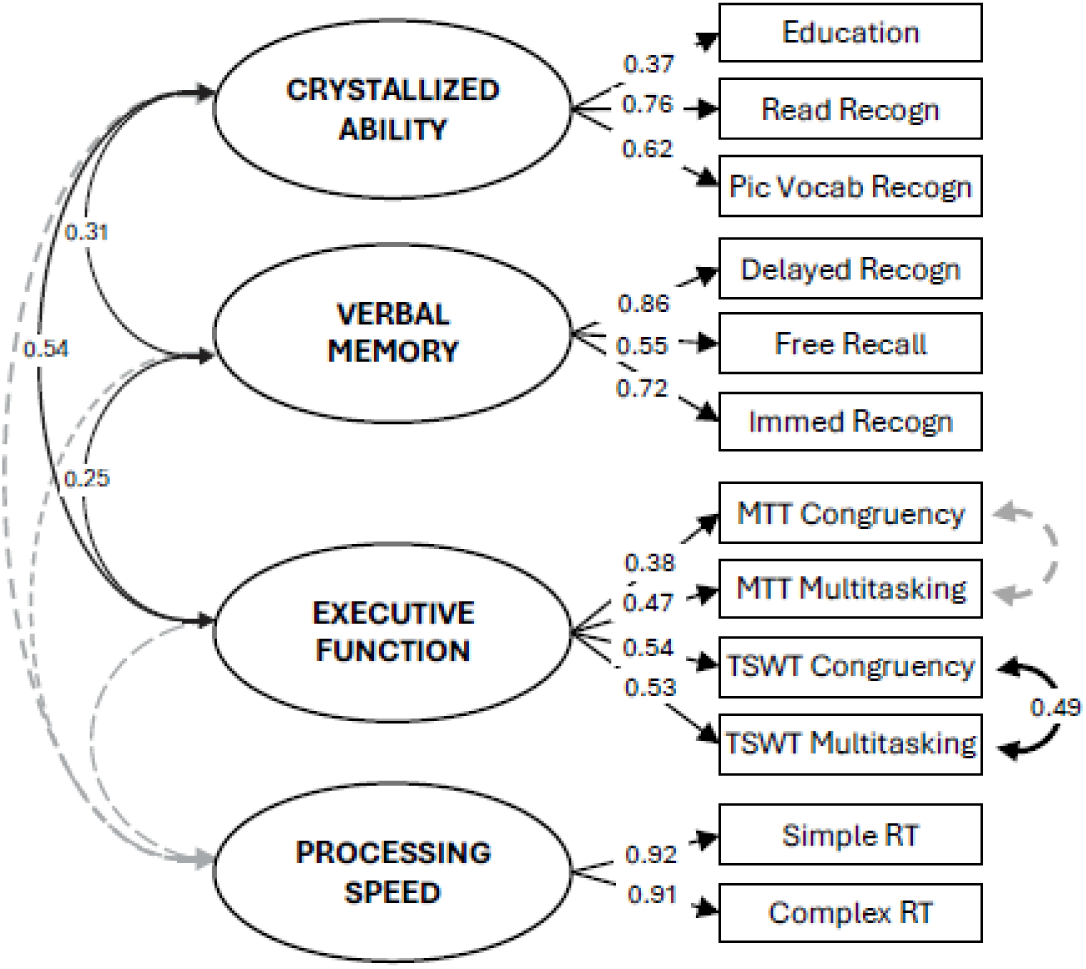
Model 1: Cognitive Ability Model. Standardized covariance coefficients (in 2-way arrows) show associations between latent cognitive factors (ovals). Standardized factor loadings of latent factors on manifest variables (rectangles) are shown in 1-way arrows. Note: For all models, where necessary, manifest variables have been reverse scored, so that for both latent and manifest variables, a high score represents good level (e.g., high education, good recall, low cost, fast RT). Solid black lines indicate *p* < 0.05; dashed grey lines indicate *p* ≥ 0.05. For cognitive variable names, see Table 1. For confidence intervals and p-values, see: https://osf.io/pu8st/.

Crystallized Ability, Verbal Memory, and Executive Function factors correlated moderately to strongly with each other (*r* = 0.24-0.52, *p* < 0.05), with higher crystallized ability associated with higher executive function and better verbal memory. The Processing Speed factor was not significantly associated with the other factors, consistent with representing psychomotor speed rather than a cognitive ability (Lohman, 1994).

#### Model 2: Cardiorespiratory Fitness (Figure 3)

The Cardiorespiratory Fitness factor significantly loaded on BMI, Heart Rate and MVPA, with the latter having the highest loading. This factor was found to significantly predict both Executive Function and Processing Speed, with higher fitness associated with better performance on the executive function tasks and faster processing speed. In contrast, the Cardiorespiratory Fitness factor did not predict Crystallized Ability or Verbal Memory.

**Figure 3.**
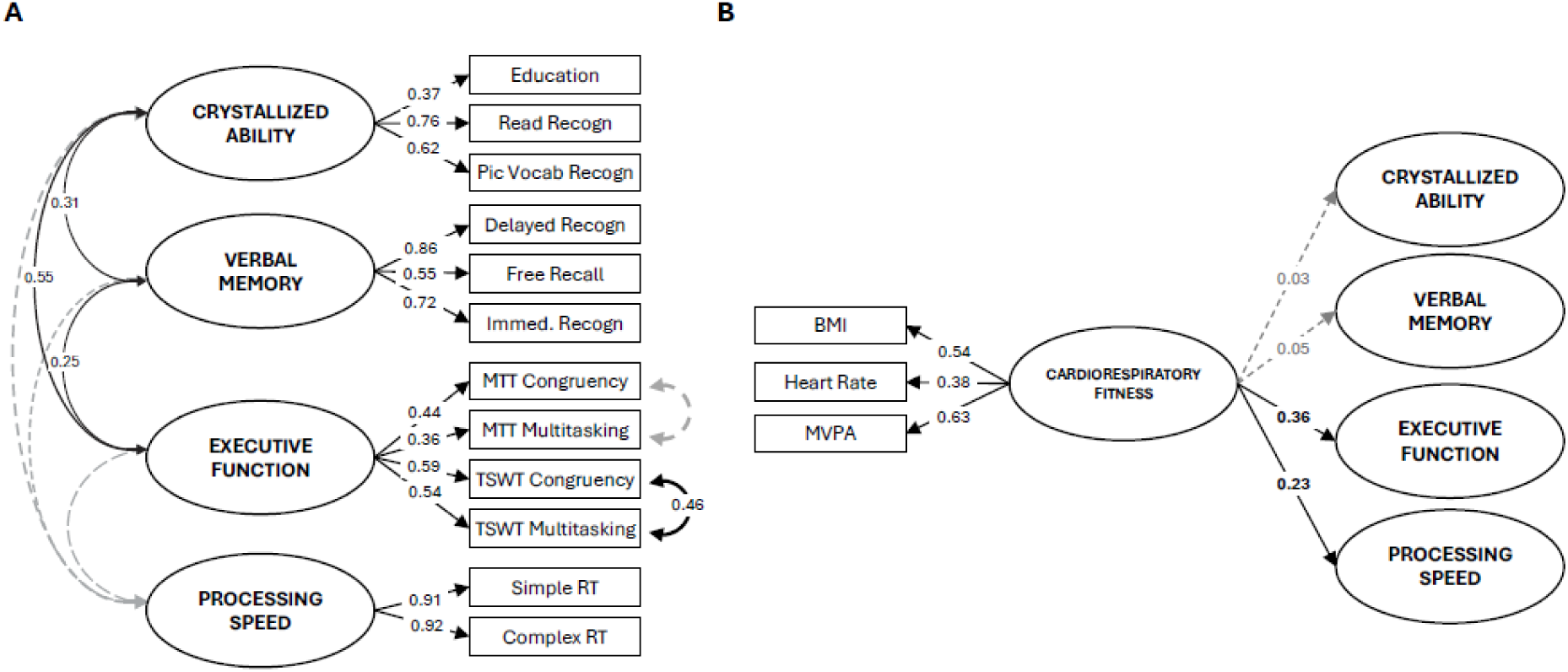
Model 2: Cardiorespiratory Fitness Model. **Panel A** shows factor loadings for cognitive factors and their covariances as estimated when examining the association between the Cardiorespiratory Fitness latent factor and each cognitive latent factor. Note that there is little difference in loadings compared to Model 1. **Panel B** shows standardized factor loadings of the Cardiorespiratory Fitness latent factor on manifest variables (*abbreviations: BMI: body mass index; MVPA: moderate-to-vigorous physical activity*) and standardized path coefficients from the Cardiorespiratory Fitness factor to each cognitive latent factor. See Figure 2 legend for all other notations.

#### Model 3: Cardiometabolic Health (Figure 4)

The Cardiometabolic factor significantly loaded on systolic and diastolic blood pressure, glucose, HDL, triglycerides, and BMI, with weakest loadings for systolic blood pressure and highest for BMI.

**Figure 4.**
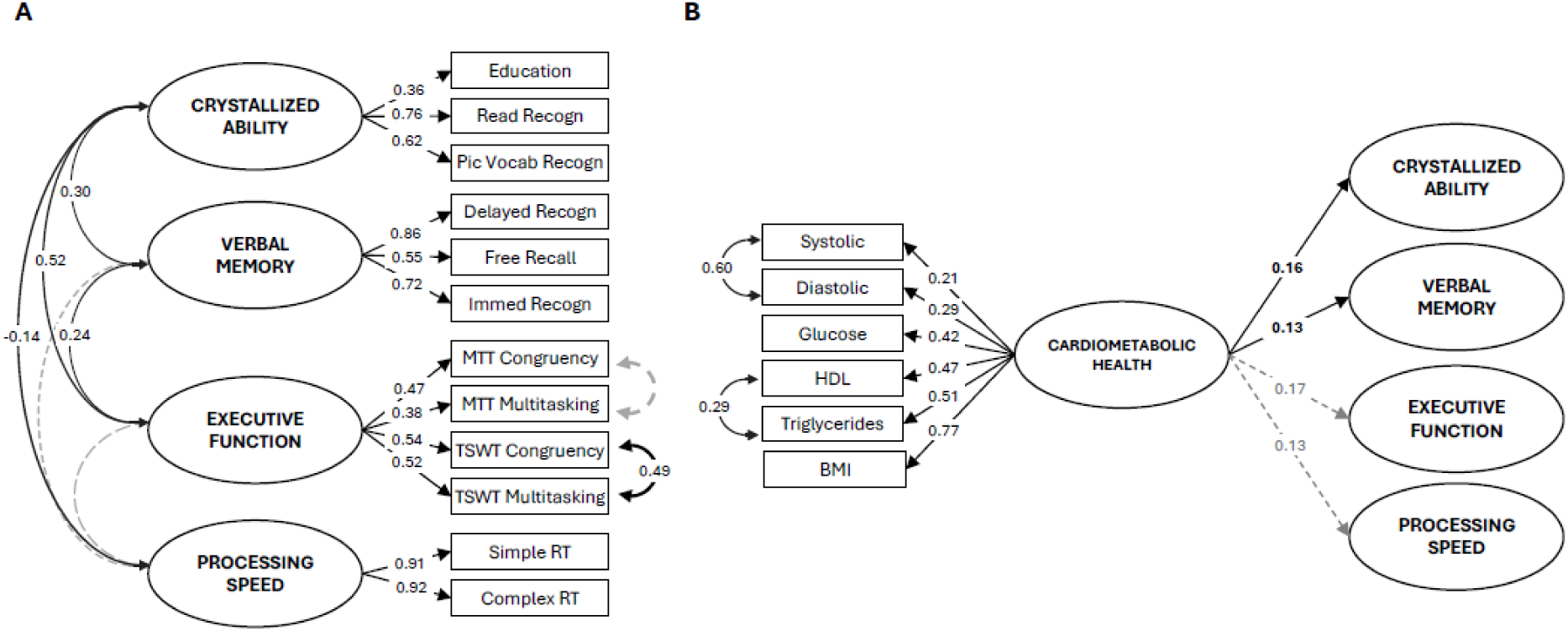
Model 3: Cardiometabolic Health Model. **Panel A** shows factor loadings for cognitive factors and their covariances as estimated when examining the association between the Cardiometabolic Health latent factor and each cognitive latent factor. Note that there is little difference in loadings compared to Model 1. **Panel B** shows standardized factor loadings of the Cardiometabolic Health latent factor on manifest variables (*abbreviations: Systolic: systolic blood pressure; Diastolic: diastolic blood pressure; HDL: high-density lipoprotein cholesterol; BMI: body mass index*) and standardized path coefficients from the Cardiometabolic Health factor to each cognitive latent factor. See Figure 2 legend for all other notations.

The Cardiometabolic Health factor weakly but significantly predicted Crystallized Ability and Verbal Memory, with higher cardiometabolic health associated with higher crystallized ability and verbal memory. In contrast, Cardiometabolic Health did not predict Executive Function or Processing Speed.

#### Model 4: Cardiovascular Health: Cardiometabolic and Cardiorespiratory Fitness (Figure 5)

In this joint model, the Cardiometabolic Health factor more strongly loaded on BMI than the Cardiorespiratory Fitness factor.

**Figure 5.**
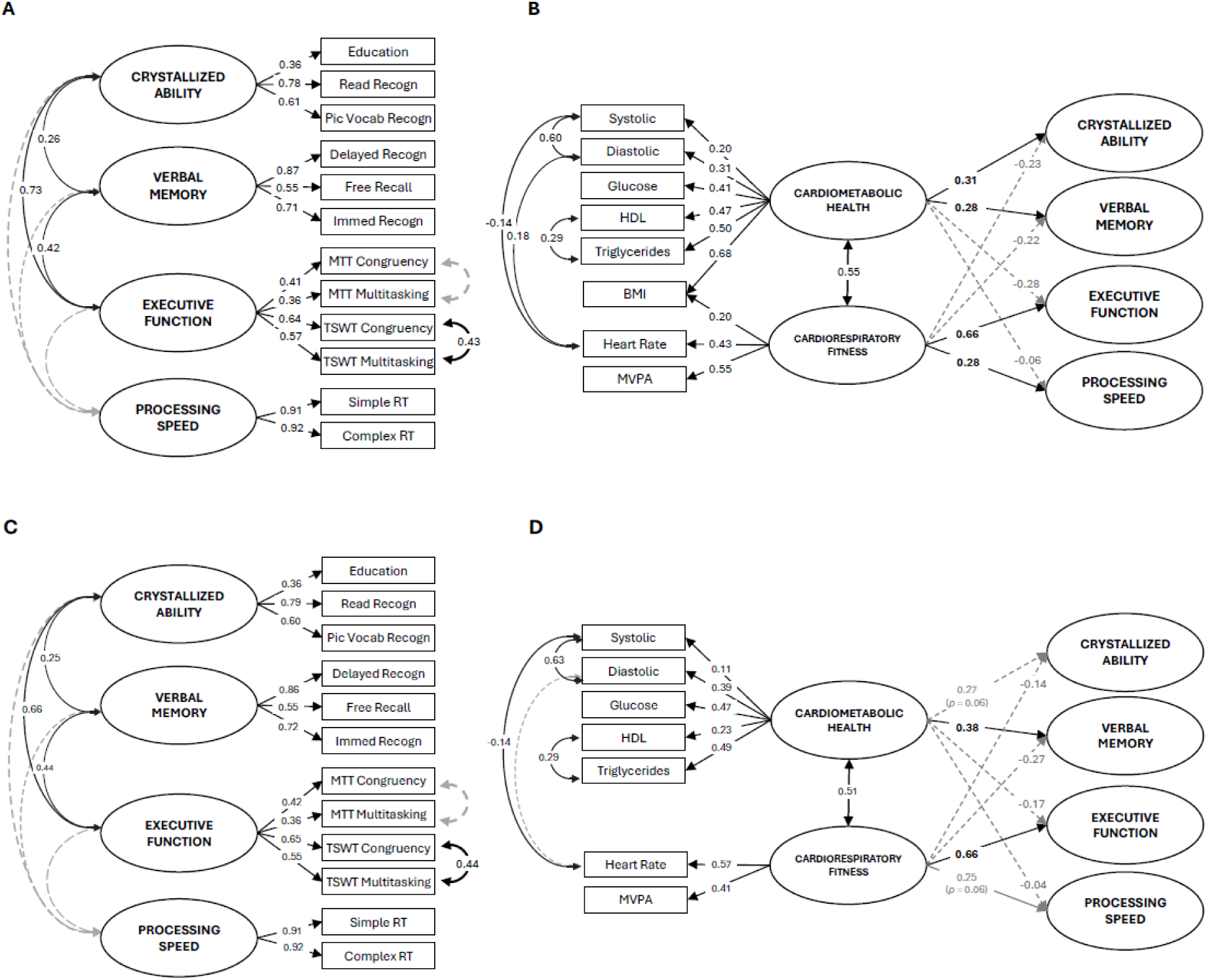
Model 4: Cardiovascular Health Model. This model includes both Cardiorespiratory Fitness and Cardiometabolic Health latent factors. It shows standardized covariance coefficients of their association, and how each latent factor relates to each cognitive factor, after controlling for their shared variance. **Panels A and B** present the model with both Cardiorespiratory Fitness and Cardiometabolic Health latent factors allowed to load on BMI. **Panels C and D** present the model without BMI. See Figure 2-4 legends for all other notations.

Cardiorespiratory Fitness and Cardiometabolic Health factors were moderately positively correlated, consistent with the close relationship between cardiometabolic health and cardiorespiratory fitness. Despite the significant relationship between Cardiorespiratory Fitness and Cardiometabolic Health factors, this joint model closely replicated the pattern of prediction of cognitive abilities seen in Models 2 and 3, with similar loadings as their respective individual models. Specifically, Cardiorespiratory Fitness predicted Executive Function and to a lesser extent Processing Speed, but did not predict Crystallized Ability and Verbal Memory. In contrast, Cardiometabolic Health moderately predicted Crystallized Ability and Verbal Memory factors but not Executive Function or Processing Speed.

To examine whether the correlation between Cardiorespiratory Fitness and Cardiometabolic Health is due to the inclusion of BMI in both factors and whether this influenced the pattern of relationships with cognitive factors, we reran the model with BMI excluded (Figure 5C and 5D). The strong correlation between Cardiorespiratory Fitness and Cardiometabolic Health persisted after removing BMI.

Notably, the exclusion of BMI changed the strength of some relationships between predictor factors and cognitive variables. Specifically, Cardiometabolic Health most strongly predicted Verbal Memory, but only marginally predicted Crystallized Ability (*p* < .06). Similarly, Cardiorespiratory Fitness still predicted Executive Function, but only marginally predicted Processing Speed (*p* < .06).

### Sex Effects

A significant effect of sex was found for both Cardiorespiratory Fitness and Cardiometabolic Health factors [*t*(343) = 2.38, *p* = 0.01 and *t*(343) = -2.07, *p* = 0.03, respectively]. Females had significantly lower Cardiorespiratory Fitness but better Cardiometabolic Health than males.

Sex also moderated the effects of Cardiorespiratory Fitness on Processing Speed [*β* = 0.13, *F*(3,341) = 13.60, *p* < 0.001]. Specifically, while higher Cardiorespiratory Fitness was associated with faster Processing Speed in both sexes, the relationship was only significant in females (*p* < 0.001; Males: *p* = 0.08). No other significant moderating effects of sex were found.

## 4. DISCUSSION

In this study, we examined associations of cardiometabolic health and cardiorespiratory fitness with cognitive abilities in a large cohort of cognitively healthy, highly active older adults aged 60-70yrs. Using a series of structural equation models, we statistically distinguished the effects of cardiometabolic health and cardiorespiratory fitness on four latent cognitive abilities, after accounting for their common variance. We selected domains that are differentially impacted by aging: crystallized abilities that improve or remain largely stable at least into late 60s ((Tucker-Drob et al., 2022)), and verbal memory, executive function, processing speed that show increasing rates of decline (Harada et al., 2013). Previous studies have examined the associations between these cardiovascular factors and cognitive ability in older adults (España-Irla et al., 2021; King et al., 2023; Oberlin et al., 2025). Here we examine their distinct and shared contribution to different cognitive domains.

Better cardiometabolic health and cardiorespiratory fitness were both associated with higher cognitive performance in this age-restricted, cognitively healthy older cohort, consistent with existing literature (Flicker, 2010; Leritz et al., 2011; Yaffe et al., 2014). However, they not only showed a distinct pattern of association with different cognitive domains, but evidence consistent with a double dissociation across cognitive domains that remain relatively stable or show little decline with increasing age, and those that are highly sensitive to age-related decline. Specifically, the Cardiorespiratory Fitness factor was significantly associated with Executive Function and Processing Speed factors, but *not* Verbal Memory and Crystallized Ability factors. In contrast, the Cardiometabolic Health factor significantly predicted scores on Crystallized Ability and Verbal Memory factors, but *not* Executive Function and Processing Speed factors. These relationships were observed when each latent cardiovascular factor was modelled independently and persisted, indeed strengthened, in the combined model that accounted for their shared variance. This is the first study to report a distinct pattern of effects, and indeed evidence consistent with a double dissociation of cardiorespiratory fitness and cardiometabolic health on distinct cognitive domains.

### 4.1 Cardiorespiratory Fitness associated with Executive Function and Processing Speed

The strongest association was found between cardiorespiratory fitness and executive function. This relationship was significant across all models and persisted even when the Cardiorespiratory Fitness factor included only heart rate and level of engagement in MVPA. Similar associations have been consistently reported previously (Carson Smith et al., 2024; Erickson et al., 2022; Festa et al., 2023).

Maintaining cardiorespiratory fitness preserves cerebral blood flow and vessel elasticity, slowing or even reversing age-related deterioration in metabolically demanding brain networks and hub regions via metabolic and trophic support (Vecchio et al., 2018; Zimmerman et al., 2014). These effects are particularly strong in frontal brain regions that support executive functioning and are highly sensitive to vascular and metabolic changes common in older adults (Kong et al., 2020; Raz et al., 2007). For example, higher cardiorespiratory fitness is consistently associated with higher brain volume in frontal and temporal regions (Colcombe et al., 2006; d’Arbeloff, 2020; Hayes et al., 2013), measures of white matter microstructural organization (e.g., fractional anisotropy, radial diffusivity, and more recently, myelin water fraction; d’Arbeloff et al., 2021; Faulkner et al., 2024; Strömmer et al., 2020; Voss et al., 2013). Additionally, cardiorespiratory fitness modulates blood flow and oxygenation in the prefrontal cortex more than in other brain regions, with oxygenation levels mediating part of the relationship between cardiorespiratory fitness and executive function in high-fit older adults (Dupuy et al., 2015; Mekari et al., 2019; Salzman et al., 2022).

An association between cardiorespiratory fitness and processing speed has also been reported, albeit less frequently (Burzynska et al., 2020; Oberlin et al., 2025; Orland et al., 2021; Pentikäinen et al., 2019). The mechanisms underlying this relationship are likely to be similar to those discussed above for executive functions. For example, higher cardiorespiratory fitness may slow the progression of age-related vascular changes, thereby protecting white matter microstructural organization (Johnson et al., 2020), both of which have been shown to predict performance on various psychomotor tasks (Hoagey et al., 2021; Kerchner et al., 2012; Rabin et al., 2019; van den Heuvel et al., 2006).

Despite well-established links between cardiorespiratory fitness, hippocampal structure, and memory performance (Erickson et al., 2011), in this age-restricted older group, we did not find an association between cardiorespiratory fitness and verbal memory performance. This may be due, at least partly, to the specific memory task used here. The CANTAB Verbal Recognition Memory Task assesses immediate word recall and immediate and short-delay recognition memory. These processes rely heavily on short-term memory and less heavily on episodic and working memory, domains more susceptible to age-related decline (Nyberg and Pudas, 2019). Bullock et al., (2018) found fitness-related benefits only in high-interference episodic memory tasks, which place greater demands on working memory and memory-related inhibition, processes particularly vulnerable to age-related decline (Biss et al., 2013; Hasher and Zacks, 1988).

More broadly, there is weak and inconsistent evidence specifically linking cardiorespiratory fitness to verbal memory. A systematic review highlighted the context-dependent nature of fitness-related memory benefits (Rigdon and Loprinzi, 2019). For instance, Dougherty et al. (2017) showed fitness effects on verbal memory only in males, while Schultz et al. (2015) reported associations between fitness and verbal episodic memory exclusively in older adults with high amyloid-beta (Aβ) burden. Longitudinal evidence also suggests that the effects of cardiorespiratory fitness on memory may not be as robust as its effects on executive functions and processing speed. Pentikäinen et al. (2019) and Pinto Pereira et al. (2023) found that cardiorespiratory fitness was more consistently associated with measures of executive function and processing speed than with verbal memory after adjusting for confounding variables such as age, sex, and education. Together, these findings suggest that the relationship between cardiorespiratory fitness and memory, particularly verbal memory, may be weak and more cohort-dependent compared to its effects on other cognitive domains.

### 4.2 Cardiometabolic Health associated with Crystallized Ability and Verbal Memory

The Cardiometabolic Health factor (i.e., comprised primarily of blood pressure and blood-based metabolic biomarkers) loaded consistently on Crystallized Ability and Verbal Memory factors, and these associations were stronger when accounting for Cardiorespiratory Fitness (i.e., β = 0.13-0.16 to 0.28-0.31).

Cardiometabolic health was not significantly associated with either executive functions or processing speed, cognitive domains that are more sensitive to aging and cardiovascular health than crystalized ability (Bucur and Madden, 2010). Therefore, the relationship between cardiometabolic health and crystalized ability, in the absence of a relationship with executive functions and processing speed, is unlikely to reflect progressive deterioration of brain vascular or cortical structure.

An alternative interpretation is that this effect arises from variability in socioeconomic status and/or educational attainment. The Crystallized Ability factor loaded on years of formal education and two verbal tests from the NIH Toolbox that capture vocabulary size and verbal proficiency - abilities widely recognized as proxies for crystallized ability (Buades-Sitjar and Duñabeitia, 2022; de Oliveira et al., 2014; Salas et al., 2021; Stern, 2002). Across the lifespan, level of cardiovascular health is lower for people from lower SES, who often have lower level of education (Paterson, 1991; Schultz et al., 2018). Low SES is also associated with poorer health literacy, which itself is associated with poorer health outcomes (Berkman et al., 2011; Svendsen et al., 2020). Years of formal education is closely linked to SES and also a significant predictor of improved cardiovascular health and associated mortality (Magnani et al., 2024; Stringhini et al., 2018).

Therefore, a plausible interpretation of the relationship between cardiometabolic health and crystalized ability is that, amongst this age-restricted, community-dwelling cohort, people with lower crystalized ability were more likely to come from lower SES backgrounds and/or have lower health literacy. This could contribute to weaker access and/or adherence to healthy lifestyle choices, and therefore poorer cardiometabolic health markers for hypertension, high cholesterol and diabetes etc.^4^

### 4.3 Cardiometabolic Health *not* associated with Executive Function and Processing Speed

Surprisingly, cardiometabolic health was associated with crystalized ability and verbal memory, but it was ***not*** significantly associated with executive functioning or processing speed. This finding goes against a strong body of work showing that these cognitive domains are highly susceptible to the presence of cardiovascular disease and risk factors, and particularly hypertension (Aljondi et al., 2020; Debette et al., 2011; Iadecola et al., 2016).

Interestingly, the Cardiometabolic Health factor loaded more on metabolic measures (i.e., lipid profiles, glucose; e.g., β = .41-.50 in Model 4) than blood pressure measures (β = .20-.31), with systolic blood pressure having the lowest value. The link between blood metabolic markers and executive function ability amongst older adults is weaker than for hemodynamic measures (i.e., systolic blood pressure; Kirvalidze et al., 2022; Peters et al., 2021). Despite blood glucose levels having the strongest load on the Cardiometabolic Health factor, the cohort’s glucose levels were well within the normal range (Table 2). So, while Liu et al., 2024 found that blood glucose level was associated with executive functions in non-diabetic older, their participants had much higher glucose levels than our cohort.

The largely healthy blood pressure range in our sample could also explain why these variables contributed relatively weakly to the Cardiometabolic Health latent factor, despite the fact that almost 50% met our operational definition of a hypertension diagnosis and over half of them were being medicated indicating a clinical diagnosis. While the effects of antihypertensive medications on current and future cognitive outcomes are still not clearly established, there is some evidence that pharmacological antihypertensive therapies are associated with reduced dementia risk and selective improvements in attention and executive function (Hajjar et al., 2012; Semplicini et al., 2011).

Another, and possibly more plausible, explanation for the above unexpected finding may be that our highly active, cognitively healthy age-restricted cohort did not have sufficiently high levels of cardiovascular risk factors for variability in cardiometabolic health to impact higher-order cognitive processes and processing speed (rather than be due to variability in SES and education, as discussed in section 4.2). As shown in Table 2, our cohort of almost 350 participants scored, on average, within the healthy range for levels of glucose, triglycerides, resting heart rate, HDL and systolic blood pressure (with some sex differences for the latter, but still within or close to healthy cutoff), below cutoff for obesity and well above recommended weekly level of MVPA. Despite the associations between cardiometabolic health and crystalized ability, it is also noticeable that, on average, the cohort was highly educated, had 7 points higher SES than the New South Wales mean, and total cognitive ability score well above established cut-offs for clinical concern. An important consideration is that recruitment into this study occurred during the COVID-19 pandemic. It is, therefore, possible that older people volunteering for research involving multiple visits to hospital and university facilities did not consider themselves to belong to highly vulnerable groups (including cardiovascular health).

### 4.3 Limitations

As noted above, when comparing this pattern of findings against other studies, it is important to recall that this cohort was on the higher end of the health scale. Therefore, our data are more indicative of variability related to preservation of function amongst high functioning older adults, than aging-related physical or cognitive decline. It is also important to note that, in this paper, we did not consider the potential contribution of other health factors (e.g., diet, sleep quality) that could impact the relationships between cardiovascular health and cognitive outcomes.

Using regression-based techniques (i.e., structural equation modeling) in a cross-sectional design limits the ability to make causal inferences (e.g., lead-lag relationships). That is, as both latent constructs were derived from physiological measures at the same time as cognitive measures, the findings cannot speak to how cumulative exposure and change over time influence cognitive ability. For example, while lower cardiorespiratory fitness and cardiometabolic health may contribute to cognitive decline, it is also plausible that the relationship is partially bidirectional (Maasakkers et al., 2021), such that cognitive decline may lead to increased sedentary behaviors, thereby reducing fitness levels and increasing cardiometabolic risk in previously high-fit, low-risk individuals. Further, SEM describes linear relationships, whereas some of the relationships associated with cognitive aging may be non-linear. This may be particularly the case for those involving vascular health conditions, such as arteriosclerosis (Bowie et al., 2024).

### 4.4 Conclusion and Future Directions

This is the first study to show evidence consistent with a double dissociation between the effects of cardiorespiratory fitness and cardiometabolic health on different cognitive domains in a large cross-sectional, age-restricted and high functioning cohort. Specifically, cardiorespiratory fitness was strongly associated with executive functioning and processing speed but not with crystallized ability or verbal memory; in contrast, cardiometabolic health was strongly associated with crystallized ability and verbal memory but not executive function and processing speed. We interpret the absence of a relationship between cardiometabolic health and tasks loading more heavily on fluid abilities as being, at least partially, related to the fact that our cohort was, on average, both cognitively and cardiometabolically healthy.

Importantly, even in this cohort, there was a strong association between cardiorespiratory fitness and both executive functioning and processing speed. These findings are supportive of a healthy aging framework that targets improvements in health literacy to maintain cardiometabolic health across the adult lifespan, while also facilitating access to lifestyle behaviors that enhance cardiorespiratory fitness and maintain functional capacity in old age.

## Footnotes

1 Also referred to as cognitive control (e.g., Gratton et al., 2018) or attentional control (e.g., von Bastian et al., 2020).

2 In SEM, when selecting manifest variables for a latent variable it is important to ensure that the factor captures the intended theoretical construct rather than being disproportionately influenced by a subset of indicators. While our dataset includes an additional measure of body composition (i.e., waist-hip ratio, alongside BMI) and other lipid profile markers (i.e., total cholesterol and LDL, in addition to HDL), we deliberately excluded these variables from modeling. Including them would result in the latent variables being overly dominated by highly correlated indicators shifting their meaning primarily toward body composition/lipid profile. To maintain conceptual clarity, we selected variables that are both widely used in the literature (BMI instead of waist-hip ratio; Czernichow et al., 2011; Huxley et al., 2010) and considered more reliable indicators of lipid metabolism (i.e., HDL instead of total cholesterol or LDL). Importantly, the pattern of results remains virtually unchanged when alternative indicators are used, suggesting that our choice does not bias the findings but rather enhances the interpretability of the latent variables.

3 We define sex as self-reported binary classification (i.e., female or male) consistent with assignment at birth (Heidari et al., 2016).

4 Our measure of SES (i.e., IRSAD) is based on postcode of residence, rather than individual measure of wealth. It is useful in characterizing the average SES of the cohort against population levels but is not sensitive enough to be used as a mediator to test the above interpretation.

## Notes

https://osf.io/pu8st/

### Competing Interest Statement

The authors have declared no competing interest.

